# Brain hubs defined in the group do not overlap with regions of high inter-individual variability

**DOI:** 10.1101/2021.10.17.464723

**Authors:** D.M. Smith, B. T. Kraus, A. Dworetsky, E. M. Gordon, C. Gratton

## Abstract

Connector ‘hubs’ are brain regions with links to multiple networks. These regions are hypothesized to play a critical role in brain function. While hubs are often identified based on group-average functional magnetic resonance imaging (fMRI) data, there is considerable inter-subject variation in the functional connectivity profiles of the brain, especially in association regions where hubs tend to be located. Here we investigated how group hubs are related to locations of inter-individual variability, to better understand if hubs are (a) relatively conserved across people, (b) locations with malleable connectivity, leading individuals to show variable hub profiles, or (c) artifacts arising from cross-person variation. To answer this question, we compared the locations of hubs and regions of strong idiosyncratic functional connectivity (“variants”) in both the Midnight Scan Club and Human Connectome Project datasets. Group hubs defined based on the participation coefficient did not overlap strongly with variants. These hubs have relatively strong similarity across participants and consistent cross-network profiles. Consistency across participants was further improved when participation coefficient hubs were allowed to shift slightly in local position. Thus, our results demonstrate that group hubs defined with the participation coefficient are generally consistent across people, suggesting they may represent conserved cross-network bridges. More caution is warranted with alternative hub measures, such as community density, which are based on spatial proximity and show higher correspondence to locations of individual variability.

## 1. Introduction

In the past two decades there has been a steady increase in the application of network science methods to cognitive neuroscience, particularly in the analysis of functional networks measured with fMRI. Functional brain networks are sets of brain regions with inter-correlated fMRI Blood Oxygen Level Dependent (BOLD) signals. These networks are present when subjects are engaged in a task or at rest (Gratton et al., 2018a). Different functional brain networks have been implicated in distinct psychological functions including sensory, motor, memory, self-referential processing, and cognitive control (Biswal et al., 1995; Dosenbach et al., 2007; Dosenbach et al., 2006; Greicius et al., 2003; Seeley et al., 2007; Thomas Yeo et al., 2011). However, many complex tasks require integration of distinct functional systems, requiring an understanding of the interactions between brain networks (Bullmore and Sporns, 2009; Gratton et al., 2018b; Sporns, 2010). Network science methods provide an opportunity to quantitatively assess the distributed interactions within and across these diverse networks, as well as the role of specific regions within this network structure.

Connector hubs (from this point forward simply referred to as hubs^1^) are specialized nodes within a complex system that have connections distributed across networks (Guimerà and Nunes Amaral, 2005; Power et al., 2013). Hubs appear to have an important role in brain networks, just as in many other complex systems (Bullmore and Sporns, 2009; Sporns, 2010; Van Den Heuvel and Sporns, 2011). Their position between different networks suggests that hubs may play an integrative role in brain function, perhaps associated with linking the distinct processes associated with different networks (Bertolero et al., 2018; Bertolero et al., 2017; Gratton et al., 2018b). Evidence in favor of this view comes from studies showing that hub activity is linked to a variety of tasks and cognitive processes (Bertolero et al., 2015; Cole et al., 2013) and their functional connectivity varies across task contexts (Cole et al., 2013; Gratton et al., 2016). Brain lesion literature also suggests that hubs play a critical role in network organization and brain function. Lesions to hubs lead to wide-spread cognitive deficits, relative to lesions to non-hub regions (Warren et al., 2014) and damage to hubs is has been associated with decreased brain network segregation (Gratton et al., 2012). Jointly, these findings suggest that hubs may play a central role in facilitating inter-network communication that enables various complex behaviors.

However, most studies of functional networks and hubs have been conducted on group data, averaged across participants. The last few years have seen a substantial growth in studies showing that there is a considerable amount of inter-subject variation in functional connectivity (Bijsterbosch et al., 2018; Finn et al., 2015; Gordon et al., 2017a; Gordon et al., 2017b; Gratton et al., 2018a; Kong et al., 2019; Miranda-Dominguez et al., 2014; Mueller et al., 2013; Seitzman et al., 2019). Reliance on group average data may obscure important individual differences (Smith et al., 2021), including variation in hub regions (Gordon et al., 2018). When a large amount of high-quality fMRI data is collected, reliable individual-level network maps can be obtained (Gordon et al., 2017b; Laumann et al., 2015). While individually-mapped networks generally resemble those found in group average data, in each participant particular locations show especially high deviation from the group average (Seitzman et al., 2019). These regions of idiosyncratic connectivity are called “network variants”. Variant regions have been shown to be stable within individuals, exhibit activation patterns similar to the network they connect with, and relate to behavioral measures collected outside the scanner (Seitzman et al., 2019). These findings suggest that variants are trait-like features of brain organization.

Locations of high inter-individual variability are most prominent in higher-level association regions (Finn et al., 2015; Gratton et al., 2018a; Kong et al., 2019; Mueller et al., 2013), with network variants most frequently found in the lateral frontal cortex and near the temporal-parieto-occipital cortex (Seitzman et al., 2019). Hubs are also typically found in association regions, especially the frontoparietal and cinguloopercular “control” networks (Cole et al., 2013; Power et al., 2013). Thus, an important question is how hubs relate to locations of individual variation in functional connectivity and more specifically are hubs defined at the group level driven by inter-subject connectivity profile variation.

Hubs, as previously measured in large groups (“typical” hub locations), and inter-individual variability in functional connectivity could be related in at least three different ways. One possibility is suggested by the vital role that hubs seem to play in tasks (Bertolero et al., 2018; Bertolero et al., 2015; Gratton et al., 2016; Warren et al., 2014), and the negative impact of damage to these regions (Gratton et al., 2012; Warren et al., 2014). This view would support the idea that hubs are critical brain locations that should be conserved across individuals, with major variations causing a significant negative impact on brain function and cognition, in the same way that nearly all humans are born with two functioning lungs If so, we would predict that, despite the high concentration of hubs in association regions, hubs will not overlap with locations of strong inter-individual variation.

A second, contrasting, hypothesis is that, as hubs are locations with variable functional connectivity across networks (and task contexts (Bertolero et al., 2018; Cole et al., 2013; Gratton et al., 2016)), hubs may be locations with generally malleable connectivity profiles, including profiles that can differ strongly across subjects. This would predict a correspondence between hub locations and locations of inter-individual variability. In this view, group hubs would still show connectivity across multiple networks in individual people, but the networks bridged by a given hub location would be variable across individuals.

Finally, it is possible that typical hubs observed in group average data are artifactual hubs, representing locations of high network variability *across* people rather than a hub (an area with network connectivity evenly distributed across multiple networks) *within* a person. That is, a group-level hub could represent an area that is coupled with a single network within each individual but vary in which network is present across individuals, yielding an average connectivity profile that has connectivity evenly distributed across multiple networks. This profile would be mistaken for a hub if researchers focus on analyzing a group-level connectivity map. These scenarios are illustrated jointly in Figure 1. Note that it is also possible that different scenarios apply to different hubs.

**Figure 1:**
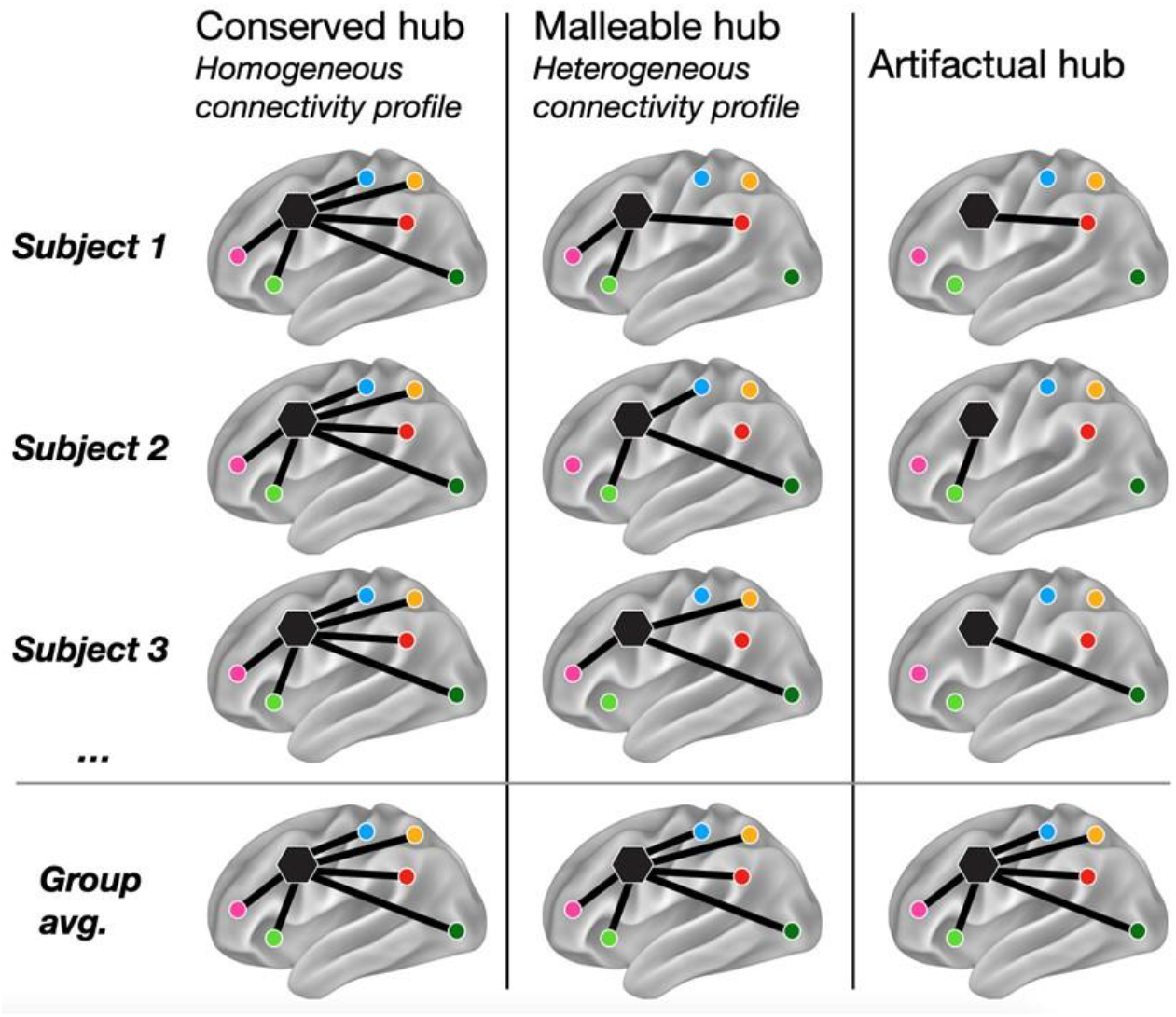
Schematic showing how group hubs may be organized within single individuals. The first column represents a “conserved hub”, a hub that has similar cross-network connectivity in each individual and in group-average data. The second column shows a “malleable hub”, a hub consistently present at the same location across individuals, but with distinct profiles in each individual (i.e., connecting to different networks). The third column represents an “artifactual hub”: within each individual, this location is a non-hub with connections to just one single network, but there is a great of inter-subject variability in the identity of this network. When data is averaged across subjects it appears that connectivity is evenly distributed across networks and makes this location appear to be a hub.

The goal of this study is to address these different possibilities by determining how group hub locations relate to areas of individual variability in functional connectivity. We examined the relationship between hubs and measures of individual variability in both a deep “precision” fMRI dataset (Midnight Scan Club; MSC, N =9 participants with ~5 hrs of resting-state fMRI) and the large Human Connectome Project dataset (HCP; N = 752, with 1 hr of resting-state fMRI). We focused primarily by examining variability at hubs defined using the common participation coefficient metric for connector hubs (Bullmore and Sporns, 2009; Guimerà and Nunes Amaral, 2005), a measure of how distributed a region’s connections are across different networks. Some additional analyses were conducted on an alternative metric of connector hubs termed “community density” (Power et al., 2013), a measure of how spatially proximal a region is to diverse networks.

## 2. Methods

### 2.1 Overview and datasets

We analyzed data from 9 highly sampled subjects from the Midnight Scan Club (Gordon et al., 2017b) and 752 subjects from the Human Connectome Project dataset (Van Essen et al., 2012). The HCP subjects are an expanded sample from that used in Seitzman et al., (2019) including all low motion individuals regardless of familial relationship (see Seitzman et al., 2019 for additional details on the composition and exclusion criteria). In each of these datasets, we identified locations of individual variability. A dataset of 120 healthy adults was used as a group average reference and to provide locations of group-average connector hubs (Power et al., 2011; Power et al., 2013). This dataset has been described in detail in Power et al (2013).

### 2.2 Data acquisition

For all three datasets collection protocols were approved by the Washington University Internal Review Board. HCP data was acquired on a custom Siemens 3T Skyra with a custom 32-channel head coil (Van Essen et al., 2012). The HCP scanning protocol included a pair of T1-weighted (256 slices, 0.7 mm^3^ isotropic resolution, TE = 2.14ms, TR = 2400ms, TI = 1000ms, flip angle = 8 degrees) and a pair of T2-weighted (256 slices, 0.7 mm^3^ isotropic resolution, TR = 3200ms, TE = 565ms) images (Glasser et al., 2013). Functional scans were collected using a multi-band sequence with MB factor 8, isotropic 2 mm^3^ voxels, TE of 33ms, and TR of 720ms (Glasser et al., 2013; Van Essen et al., 2012). One hour of resting state data was acquired per subject in 15 min. intervals over two separate sessions (Van Essen et al., 2012).

For the MSC, high-resolution T1-weighted (224 slices, 0.8 mm^3^ isotropic resolution, TE = 3.74ms, TR = 2400ms, TI = 1000ms, flip angle = 8 degrees), T2-weighted (224 slices, 0.8 mm^3^ isotropic resolution, TE = 479ms, TR = 3200ms) both with 0.8 isotropic resolution, and resting state BOLD data were collected on a Siemens 3T Magnetom Tim Trio with a 12-channel head coil (Gordon et al., 2017b). Functional scans were collected with a gradient-echo EPI sequence, isotropic 4mm^3^ voxels, TE of 27ms, and TR of 2200ms (Gordon et al., 2017b). The MSC dataset acquired 5 hours of resting state data per subject in 30 min. blocks over 10 separate sessions (Gordon et al., 2017b).

The WashU-120 dataset was collected on a Siemens MAGNETOM Tim Trio, 3T scanner with a Siemens 12 channel Head Matrix Coil. Both T1-weighted (127 slices, 1 mm^3^ isotropic resolution, TE = 3.06ms, TR = 2400ms, TI = 1000ms, flip angle = 8 degrees) and T2-weighted (32 slices, 2 × 1 × 4 mm^3^ resolution, TE = 84ms, TR = 6800ms) scans were collected (Power et al., 2013; Power et al., 2014). The amount of resting state data collected per subject ranged from 7.7 to 16.5 min (TE = 27ms, isotropic 4mm^3^ voxels; TR = 2500ms, flip angle = 90 degrees).

### 2.3 Preprocessing

#### 2.3.1 General Preprocessing

For each of the three datasets the T1-weighted images were processed via automatic segmentation of the gray matter, white matter, and ventricles in Freesurfer 5.3 (Fischl et al., 2002). The default recon-all command in Freesurfer was then applied to produce the anatomical surface for each subject (Dale, 1999). In the MSC dataset, these surfaces were manually edited to improve the quality of the registration. The surfaces were registered to the fs_LR_32k surface space via the procedure outlined in Glasser et al. (2013).

For the HCP dataset the volumetric BOLD time series from each run were concatenated. Slice timing correction was not performed for the HCP dataset in accordance with the minimal preprocessing pipeline guidelines (Glasser et al., 2013). Field inhomogeneity distortion correction was conducted using the mean field map. Motion correction was conducted using rigid body transforms aligning to the first frame of the first run. After this step whole-brain intensity values across each BOLD run were normalized to achieve a mode value of 1000 (Miezin et al., 2000). This dataset was processed in MNI atlas space with 2 mm isotropic voxels.

The preprocessing pipelines used for the MSC and WashU-120 datasets were almost identical to the HCP with some minor exceptions. Field inhomogeneity distortion correction using the mean field map was applied to all sessions for the MSC dataset but not for the WashU 120 given that field maps were not collected for this dataset (Gordon et al., 2017b; Laumann et al., 2015). Slice timing correction was performed in both the MSC and WashU-120 datasets using sinc interpolation to account for temporal misalignment in slice acquisition time. This was followed by motion correction which was performed within and across BOLD runs (aligned to the first frame of the first run) via a rigid body transformation. Then whole-brain intensity values across each BOLD run were normalized to achieve a mode value of 1000 (Miezin et al., 2000). For the WashU-120 functional BOLD data was then registered directly to a high resolution T1-weighted structural image from each participant. For the MSC functional BOLD data was first registered to a T2-weighted image and then to the T1. An affine transformation was used for registration in both datasets. The T1-weighted image was aligned to a template atlas (Lancaster et al., 2000) conforming to Talairach stereotactic atlas space using an affine transformation. All computed transformations and re-sampling to 3 mm isotropic voxels were simultaneously applied at the end of these steps

#### 2.3.2 Resting State Connectivity Pipeline

Steps were taken to mitigate the influence of artifacts on resting state BOLD time series. The impact of nuisance signals was attenuated via regression of average signal from the white matter, ventricles, global signal, motion parameters, as well as their derivatives and expansion terms (Friston et al., 1998; Power et al., 2014). The impact of motion was further mitigated via the removal of frames with framewise displacement > 0.2 mm, in addition to sequences containing less than 5 contiguous low motion frames, the first 30 seconds of each run, and runs with < 50 low motion frames (Power et al., 2014). In the HCP dataset, before censoring high-motion frames, motion parameters were low-pass filtered at 0.1 Hz to reduce the effects of respiratory artifacts on motion estimates stemming from the short-TR multi-band acquisition (Fair et al., 2020; Gratton et al., 2020; Siegel et al., 2017). Then a filtered FD threshold of 0.1 mm was applied to censor frames. The same filtering procedure was also applied to two MSC subjects (MSC03 and MSC10) with respiratory contamination in their motion parameters. In all cases, flagged head motion frames were removed and the time points were replaced with interpolated data using a power-spectral matched approach (Power et al., 2014), after which a bandpass filter (0.009 Hz-0.08 Hz) was applied to the data.

As previous results have indicated that ~45 min. of low motion data are necessary to achieve high reliability of network variant locations (Kraus et al., 2021; Seitzman et al., 2019), we then removed any participant with less than 75%, or 45 min., of data in the HCP. In the 1200-HCP release, this resulted in 752/1206 final participants. In the MSC dataset 9/10 participants were retained (Gordon et al., 2017b). The excluded MSC participant was removed due to high motion and drowsiness (Gordon et al., 2017b; Laumann et al., 2017).

For all datasets the processed BOLD data were mapped to each individual’s native midthickness surface via the ribbon-constrained sampling procedure (Marcus et al., 2013). Then, the mapped data were registered to the fsaverage surface in one step using the deformation map generated from the ribbon-constrained sampling procedure described in Glasser et al., (2013). Next, smoothing was conducted via a geodesic Gaussian smoothing kernel to the surface registered data (FWHM = 6 mm, sigma = 2.55) (Gordon et al., 2016; Marcus et al., 2011). Temporally interpolated frames were then removed prior to functional connectivity analysis. Functional connectivity was calculated as the Pearson correlation coefficient between different cortical locations, based on time series averaged across regions.

### 2.4 Regions of interest and functional brain networks

A set of 264 spherical (10 mm diameter) regions of interest from (Power et al., 2011) were used as a basis to define group-average brain hubs (Power et al., 2013). These regions divide into networks largely overlapping with the 14 canonical networks defined in (Gordon et al., 2017a): the default mode (DMN), visual, fronto-parietal (FP), dorsal attention (DAN), language (Lang), salience, cingulo-opercular (CO), somatomotor dorsal (SMd), somatomotor lateral (SMl), auditory, temporal pole (Tpole), medial temporal lobe (MTL), parietal medial (PMN), and parietooccipital (PON). Group hubs were defined from the 264 spherical regions, based on participation coefficient estimates previously published in Power et al. (2013) which were calculated from the group average of the WashU-120 dataset (see Section 2.5).

The 14 canonical networks are also defined at the cortical surface vertex level in the WashU-120 group average (Laumann et al., 2015). The cortical surface networks were used for the network profile analyses (see section 2.6.3). These networks were defined with the Infomap clustering algorithm (Rosvall and Bergstrom, 2008) which yielded data-driven functional network definitions for a range of edge density thresholds from 0.3%-5% (Gordon et al., 2017b). A group average network consensus map was derived by collapsing network definitions across thresholds.

Measures of inter-individual variation (spatial correlations between an individual and the group average and identification of network variants) were calculated at the single cortical vertex, rather than regional, level. These are described in Section 2.6.

### 2.5 Hub Definition

Hubs were defined by two approaches: participation coefficient and community density. Primary analyses focused on the locations of brain hubs defined in group average data relative to locations of individual differences. These were derived from the WashU-120 group-average dataset based on published values from Power et al., (2013).

#### 2.5.1 Participation coefficient hubs

The participation coefficient is a graph theoretic measure that captures how evenly distributed a node’s connections are across networks (Guimerà and Nunes Amaral, 2005); see Figure 2A for schematic). The participation coefficient for node *i* is defined as 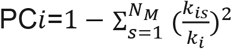, where *k_i_* is the degree (the number of edges/connections to nodes in the given node’s module/network) of node *i, k_is_* is the number of edges of node *i* to nodes in module/network *s*, and *N_M_* is the total number of networks/modules in the graph.

**Figure 2:**
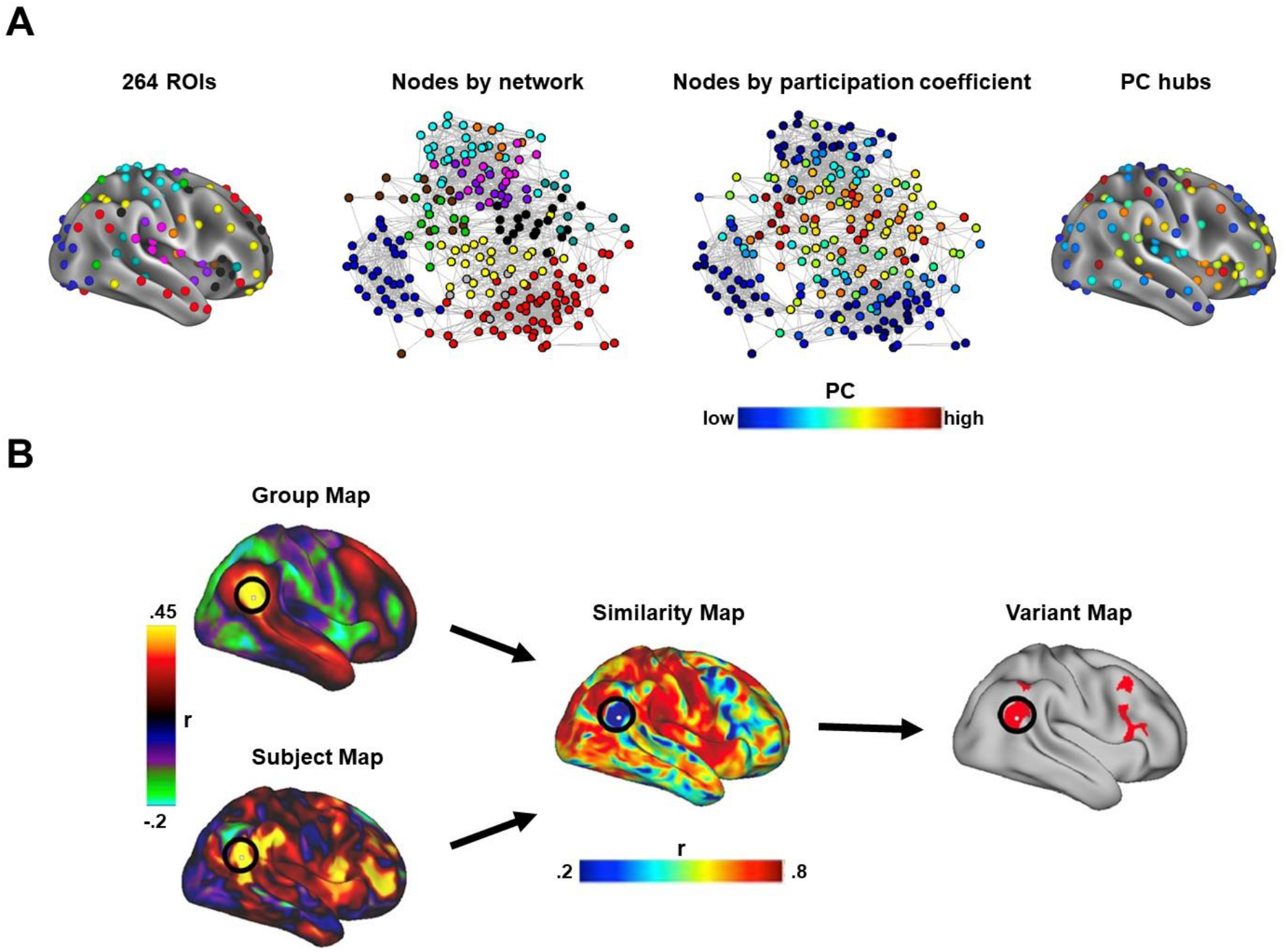
A schematic of the identification of connector hubs and locations of inter-individual variability. (A) Connector hubs are locations with connections to multiple networks. Brain network regions, or ‘nodes’ can be depicted on the brain based on their location (left image) or as a spring-embedded network (middle left), where nodes with more connections are placed closer together (colors = networks). In this depiction, it becomes clear that some nodes lie intermediate to multiple networks, a characteristic that can be quantified with the participation coefficient (Guimera & Ameral 2005; middle right). Here, we ask how connector hubs relate to locations of inter-individual variability. (B) The identification of locations of inter-individual variability starts by comparing the connectivity profile for each vertex in an individual level seed connectivity map (subject map) to the seed connectivity map for the corresponding vertex in the group average reference dataset (group map). For each vertex the spatial correlation between these two maps is calculated to produce a similarity map that represents how similar an individual is to the group at each vertex. Variants are defined as sets of at least 50 contiguous vertices, not falling in low SNR locations, that are all in the bottom similarity decile (lowest 10% of locations).

Participation coefficient hubs in the group were identified based on a previously published analysis of the large WashU-120 group-average dataset (Power et al., 2013). In brief, in that work the participation coefficient was calculated for each of 264 regions (nodes) for a range of sparsity thresholds from 2-10% edge density in 1% steps (network definitions were derived at each threshold for these calculations as well using the Infomap community detection algorithm [(Rosvall and Bergstrom, 2008)]). Participation coefficient values were then normalized and summed across thresholds to result in a final value for each region. The ten regions with the highest participation coefficient values out of a set of 264 regions of interest were selected for analysis. The MNI coordinates of the centers of each of these ten spherical regions of interest were projected to a vertex on the Conte69 midthickness surface and dilated to a 5mm radius. In addition, these locations were checked to determine if more than 30% of their vertices overlapped with a low signal mask (mean BOLD signal less than 750 computed as in (Ojemann et al., 1997)). No hub vertices overlapped with the low signal mask.

#### 2.5.2 Community density hubs

Hubs defined by community density were based on the top 10 community density peaks listed in Power et al., (2013). In brief, these were defined through the following procedure. Cortical voxels were assigned to networks using the Infomap clustering algorithm (Rosvall and Bergstrom, 2008) at a range of density thresholds (0.5-2.5% at 0.5% intervals). Community density was defined as the number of networks appearing within a radius of a given voxel (with radii ranging from 5-10mm in increments of 1mm). Values were summed across thresholds and radii after normalizing the values within each analysis, resulting in one final community density value per voxel. The MNI coordinates of the top 10 community density peaks were then projected to the cortical surface and dilated 5mm. As before, regions were checked for overlap with a low signal mask; none of these regions exceeded the overlap threshold.

#### 2.5.3 Locations of inter-subject variation in functional connectivity

The similarity of an individual’s connectivity profile and the group average connectivity profile was gauged via a spatial correlation following previously published methods (Laumann et al., 2015; Seitzman et al., 2019) (see Figure 2b for schematic). For each vertex on the cortical surface, its BOLD time series was correlated with the time series for every other vertex to form a seed correlation map. Each seed correlation map (connectivity profile for a given location) was Fisher Z transformed, vectorized, and the correlated (Pearson correlation) with the Fisher Z transformed correlation vector of the corresponding cortical vertex in the group average data map, resulting in a single similarity value for that location. Across all locations we use these values to form a map of correlations between the individual level connectivity profile and the group average connectivity profile, which we refer to in this work as a *“similarity map”*.

Areas of extreme idiosyncratic functional connectivity, which we call *“network variants”* were then defined from this map using a recently developed procedure (Seitzman et al., 2019). We began with the similarity map and then identified the locations that were most dissimilar (bottom 10%) between an individual and the group. Regions were required to be composed of at least 50 contiguous vertices, fall outside areas with a low signal (mean BOLD signal less than 750 computed as in (Ojemann et al., 1997)), and to not overlap with the network the area’s vertices were originally assigned (Seitzman et al., 2019).

In order to determine where variants are most frequently found, we created an overlap map of the network variants across participants. The frequency of variants was defined as the percentage of subjects with a variant at a given cortical location (vertex). Variant frequency maps were produced for both the HCP and MSC datasets (Seitzman et al., 2019).

### 2.6 Relationship between hubs and locations of individual variability

The relationship between hubs and locations of individual variability was analyzed in several ways. First, we examined whether the hubs collectively overlap with network variant locations (locations of extreme variability). We also examined to extent to which single hub locations overlapped with variants. Second, we used the continuous individual-to-group similarity map to examine more subtle variations at hub locations. Third, we examined the network profile of connector hubs within single individuals and employed a local spotlight procedure to determine if it was possible to slightly adjust the position of hubs to improve intersubject correspondence.

#### 2.6.1 Quantification of overlap between group-average hubs and variants

We first examined the overlap of variants with group hub locations. The top 10 group hubs were defined as described above, using both participation coefficient and community density. A variant frequency map was then produced for each dataset, capturing the frequency of variants at each cortical location. We then measured the overlap between these maps.

To determine if the frequency of variants at hub locations is greater than what would be expected for a random set of cortical locations of the same size with a similar spatial configuration, a null distribution was generated by randomly rotating a set of hubs across the cortical surface (Gordon et al., 2016). For each set of hubs (either participation coefficient or community density based) 1000 rotations were randomly generated and performed within each hemisphere. If any of the hubs intersected with the medial wall the rotation was recalculated until none of the hub locations overlapped with the medial wall. For each rotation, we calculated the average frequency of variants at rotated hubs. Across 1000 rotations, this produced a null distribution of the expected variant frequency at rotated hubs. We then compared the actual variant frequency for hubs (averaged across hub vertices) with that of the rotated distribution. This distribution was used to derive percentiles and 95% confidence intervals for the variant frequency of hubs. A similar analysis was conducted for each single hub as well, in this case comparing variation at that hub to a null made based on rotating only that hub location.

#### 2.6.2 Quantification of overlap between group-average hubs and subtle variation

The aforementioned analyses are geared towards determining if hubs overlap with areas of extreme inter-subject deviation in functional connectivity. In addition, we conducted a secondary analysis to investigate whether hubs overlapped with more subtle forms of intersubject variation. For this analysis, we used the continuous (unthresholded) individual-to-group similarity map described in section 2.5.3 for variant definition. For each vertex, a continuous value represents how similar this location is to the group average in a given MSC or HCP subject. We then determined whether this individual-to-group similarity of hubs was lower than expected by chance by comparing the values at hub locations with the values obtained through random rotations of hubs, as described above.

#### 2.6.3 Quantification of the network profile of hubs across individuals

Next, we examined the network profile of hub locations within individual participants of the MSC, to determine whether they exhibited high connectivity to multiple (similar) networks. For each participation coefficient hub, the average correlation of the hub to each of the 14 canonical networks (defined on the cortical surface; see section 2.4) was calculated. This resulted in a 14×1 network correlation vector for each participant for each hub that we call a ‘network profile’.

#### 2.6.4 Local adjustments of group hub locations

Launching from the results of the network profile procedure above (2.6.3), we asked whether group hub locations could be slightly spatially shifted within individuals to improve the correspondence of hubs across people. We used a spotlight analysis to investigate this possibility in the MSC dataset. A spotlight was formed by dilating 10mm around the central vertex of a group hub. The hub center was then moved throughout this spotlight (with hub extent still defined based on a 5mm dilation). Potential hub locations with less than 70% of its vertices within a single network were removed from consideration. These potential hub locations were then trimmed to encompass a single network (using the vertex-wise canonical network maps described in section 2.4).

At each potential hub location, we then examined how hubs were related to each of the 14 canonical networks, creating a network profile vector as described in section 2.6.3. The resulting network profiles were compared (via Euclidean distance) to a group-average reference profile based on the WashU-120 dataset. The best fitting (lowest Euclidean distance to the reference) potential hub location was selected as the final adjusted hub location for a given individual.

### 2.7 Data and code availability

All of the data have previously been made publicly available (HCP: https://www.humanconnectome.org/; MSC: https://openneuro.org/datasets/ds000224/versions/00002; WashU 120: https://legacy.openfmri.org/dataset/ds000243/). Code for analysis related to network variants in MATLAB is available at: https://github.com/GrattonLab/SeitzmanGratton-2019-PNAS; other code related to MSC processing can be found at: https://github.com/MidnightScanClub. Code related specifically to the analyses in this article will be located at this link upon publication: https://github.com/GrattonLab/.

## 3. Results

### 3.1 Overview

The aim of our investigation was to determine how group defined hubs relate to areas of variation in functional connectivity across people. We hypothesized that hubs are critical regions that should be conserved across individuals; thus they should exhibit relatively little variability across subjects. A second contrasting hypothesis is that these regions are malleable in their functional connectivity, exhibiting a high degree of variability in their connectivity profiles across participants (but still remaining a hub across participants). A final alternative is that typical hubs defined at the group level may arise from averaging non-hub regions associated with diverse single networks across individuals, creating an artifactual hub representation (Figure 1).

To test these alternatives, we examined how group hubs relate to locations of individual variation in functional connectivity. We then characterized group hubs in more detail, by examining their network profiles and determining whether small local adjustments in those profiles improved correspondence across individuals. We focused primarily on hubs defined with the often used participation coefficient, a graph theoretic metric of the distribution of a node’s connections across different networks. Secondary analyses also contrasted these results with the alternative community density metric for connector hubs, which is based on the spatial proximity of a location to multiple different networks.

### 3.2 Group hubs defined by the participation coefficient do not overlap with network variants

How do hubs vary across people? We began by examining the locations of typical group hubs, defined as the top 10 participation coefficient regions estimated from a group-average of 120 healthy young adults (*WashU-120*) in previous work (Power et al., 2013). The participation coefficient measure defines hubs as regions with distributed functional connectivity across networks (see section 2.5.1). In parallel, we identified locations of high inter-subject variability (which we term “network variants”; Seitzman et al 2019). We asked how hub locations corresponded with network variants across people.

As can be seen in Figure 3, network variant locations are especially frequent in the temporoparietal junction, lateral frontal cortex, and the dorsal posterior cingulate. Participation coefficient group hubs are also found in association systems, but more prominently in the anterior insula, superior parietal cortex, and dorsolateral and medial frontal cortex (Figure 3, Power et al., 2013). Thus, there appears to be relatively low overlap between participation coefficient hubs and locations that frequently vary across people.

**Figure 3:**
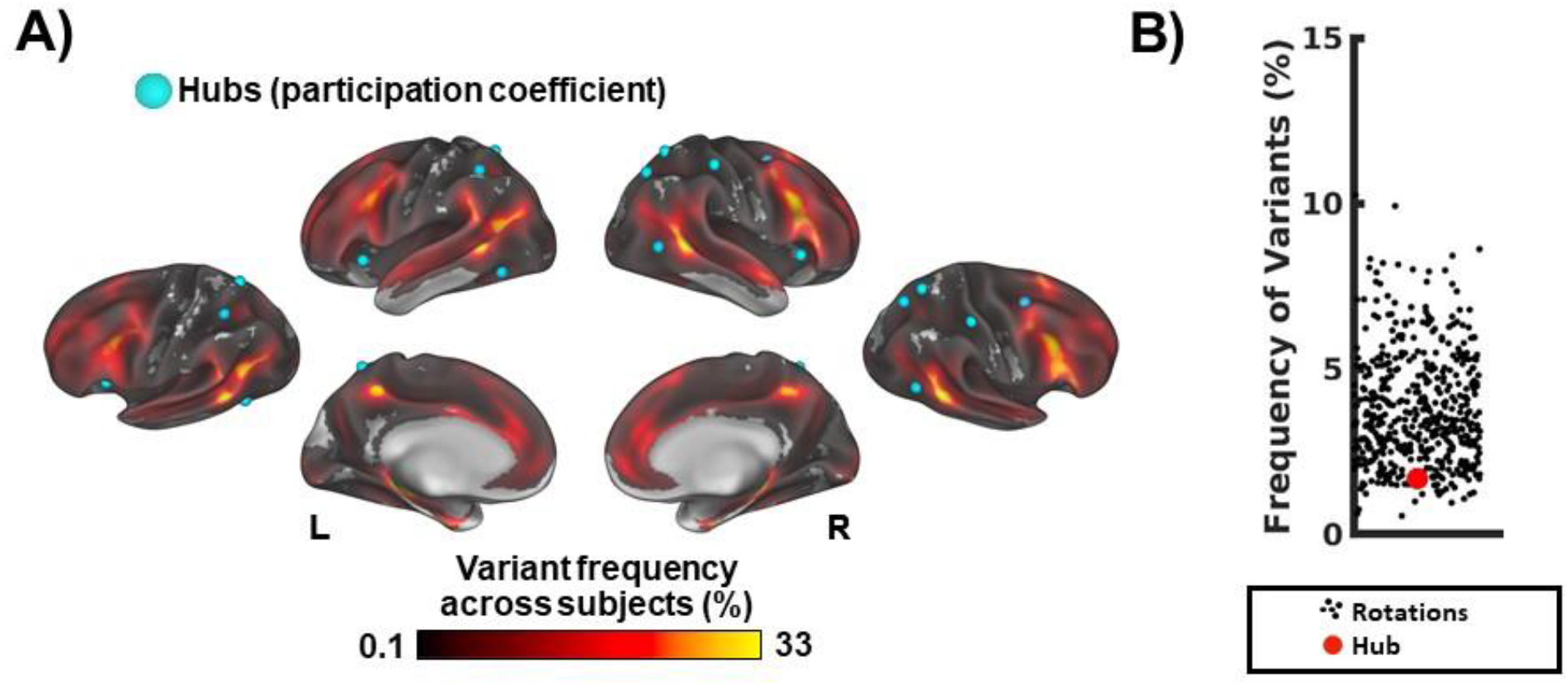
Comparison of group participation coefficient hubs to locations of inter-individual variation. A) The locations of group hubs defined using the participation coefficient are represented as light blue foci on the cortical surface. The heat map displayed on the cortical surface captures the frequency of variants (percentage of subjects with a variant at a location) based on the HCP dataset, with warmer colors indicating greater variant frequency. B) Compares the true frequency of variants at participation coefficient hubs (red dot) relative to random rotations of the hub set (black dots). Group hubs defined by the participation coefficient do not frequently overlap with areas of idiosyncratic functional connectivity.

Confirming this qualitative description, participation coefficient hubs, as a whole, occurred over variants at a low rate in the HCP dataset, within the bounds of what would be expected by chance relative to 1000 random rotations of the hub set (Fig. 3B; 1.69% of people had variants at hub locations, at the 8^th^ percentile of random rotations, 95% CI [1.22%, 7.93%]). This pattern replicated in the precision MSC dataset (Supp. Fig. 1A). Thus, group hubs defined by the participation coefficient do not show significant correspondence to areas of strong inter-subject variability.

We also examined how each single hub varied (Fig. 4). The 10 top participation coefficient hubs were separately compared to a distribution of variant frequency in the HCP. This distribution was contrasted with a null overlap distribution produced from 1000 random rotations of a region of the same size as the hubs. None of these top participation coefficient hubs deviated from what would be expected from their null distribution (frequency of network variants at single hubs: 1.59% +/- 1.55%; frequency of network variants with random rotations: left hemisphere = 3.83% +/-5.46%; right hemisphere: 3.82% +/- 5.04%). The hub with the highest overlap with network variants was in the right superior caudal portion of the frontal lobe; this location had variants in 5.47% of people, still within the bounds of what would be expected by chance (77^th^ percentile rotation). All other hubs were within a standard deviation of the null distribution’s mean.

**Figure 4:**
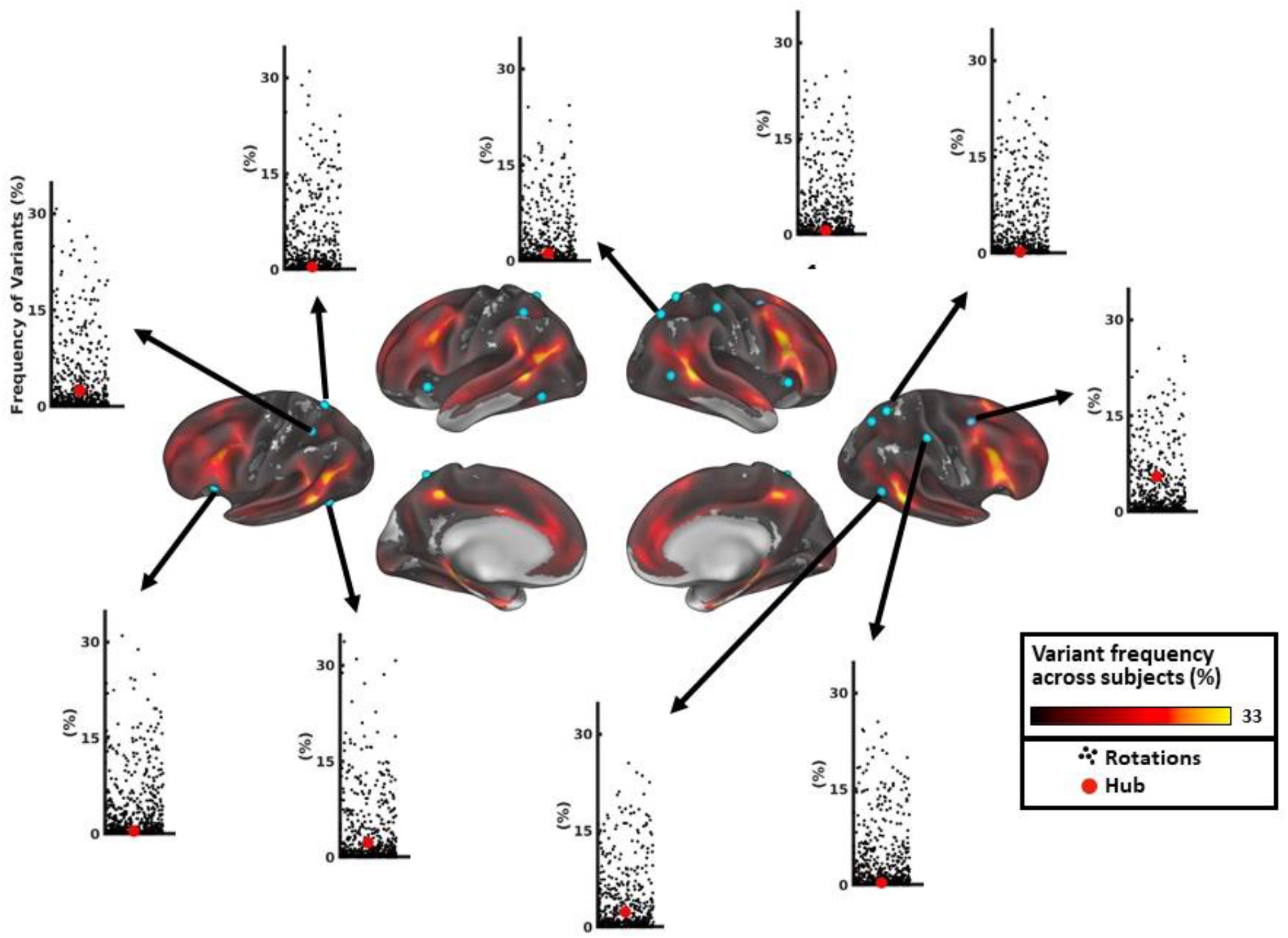
Network variant frequency at single participation coefficient hub locations. Each foci on the cortical surface represents a participation coefficient hub based on group-average data. The scatter plots capture the relationship between the true frequency of network variants at a given hub region (red dot) and the amount expected by random rotations (black dots). None of the hubs significantly differed from what would be expected from random rotations and all but one of the hubs (in the superior frontal cortex) were below the mean of the null distribution.

These findings show that group participation coefficient hubs do not frequently overlap with variants. This result is in line with the conserved hub hypothesis which states that the connectivity profiles of hubs are similar across individuals. In contrast, the malleable and artifactual hub hypotheses predict that group hubs should correspond with areas of high idiosyncratic functional connectivity.

### 3.3 Community density hubs do overlap with locations of variability

In past work, community density has been proposed as an alternative measure of connector hubs (Power et al., 2013). This measure defines hubs based on their proximity to multiple different networks, under the assumption that regions at the intersection between networks are well situated to mediate cross-network interactions. As before, we compared the locations of the top 10 group hubs, in this case defined based on community density in the same large sample of healthy young adults used in previous work (Power et al., 2013), with the map of the frequency of variants in the HCP dataset (Seitzman et al., 2019). As depicted in Figure 5A, the top community density hubs are found in somewhat similar locations to participation coefficient hubs, but more ventrally in the anterior insula, along the superior frontal cortex, and near the temporoparietal junction. As a whole, community density hubs overlapped with network variant locations significantly more frequently than what would be expected by chance, as assessed with random rotations (Fig. 5B; 10.23% of people had variants at community density hubs, at the 99^th^ percentile of random rotations, 95% CI [1.14%, 8.00%]). This result was replicated in the MSC dataset (Supp. Fig. 1B). These findings suggest that community density hubs from group-average data often overlap with areas of idiosyncratic functional connectivity, suggesting they may be identifying malleable (Fig. 1B) or artifactual (Fig. 1C) hubs.

**Figure 5:**
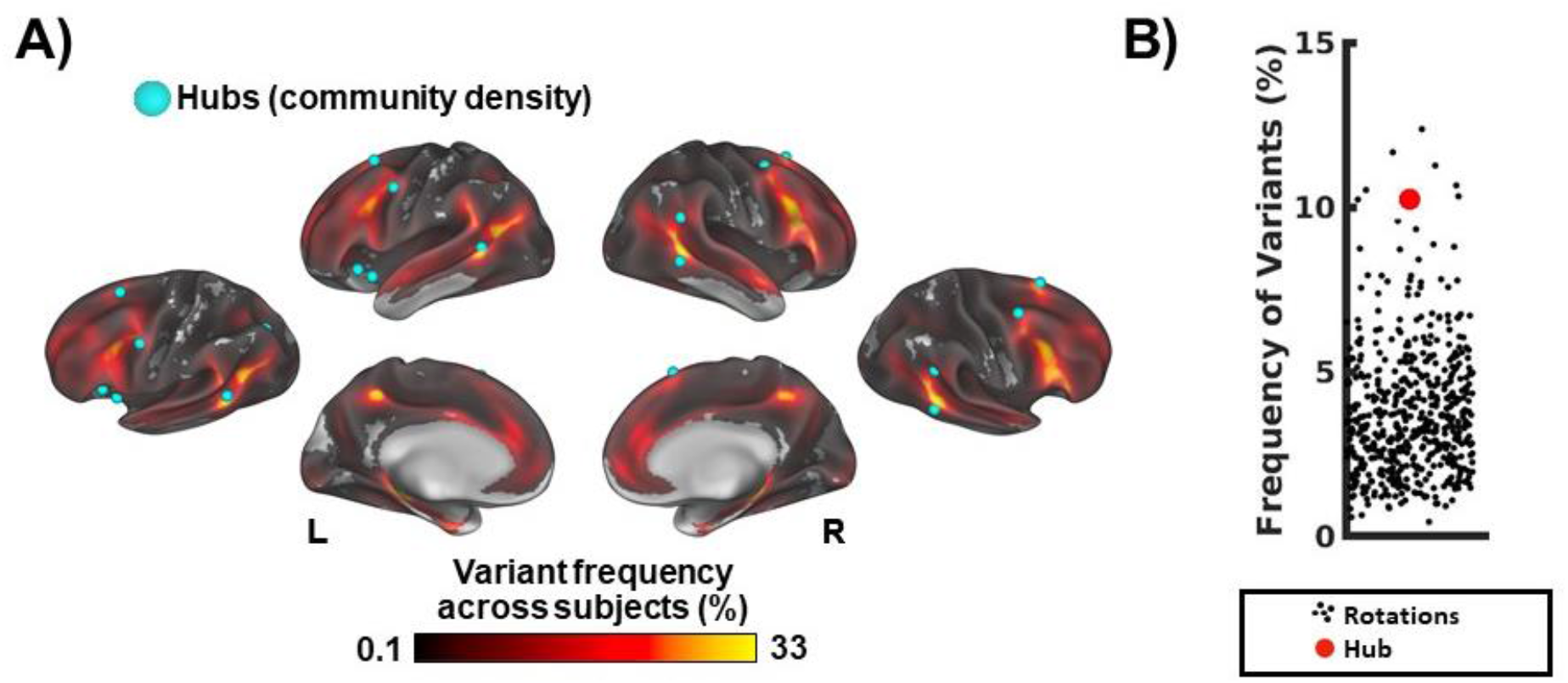
Comparison between group-average community density hubs and locations of strong inter-individual variability. A) The locations of group hubs defined using the community density metric are represented as light blue foci on the cortical surface. The heat map displayed on the cortical surface captures the frequency of network variants across people (percentage of subjects with a variant in at a location) based on the HCP dataset, with warmer colors indicating greater variant density. B) A scatter plot comparing the true frequency of variants (red dot) for the community density hub set to random rotations of the hub set (black dots). The high degree of correspondence between variants and community density hubs suggests that these hubs might be malleable or even artifactual hubs.

As before, we examined these results in more detail by quantifying the overlap of specific community density hubs with network variants in the HCP dataset (Supp Fig. 2). On average, 9.61% of people had variants over a connector hub (standard deviation: 6.44%; range: 0.58%-20.44%). Two hubs fell outside of the 95% confidence intervals of what would be expected by chance based on random rotations: one in the right superior frontal cortex and one near the left temporoparietal junction, but neither case survived FDR correction (Benjamini and Hochberg, 1995). Hubs in the left lateral frontal cortex, right temporoparietal junction, and the right lateral frontal cortex also exhibited relatively high frequency of variants. Thus, community density hubs generally have a high tendency to overlap with network variants, especially hubs in superior frontal cortex and near the temporoparietal junction.

### 3.4 Participation coefficient hubs exhibit similarity to the group-average

The previous analyses demonstrate that group-average participation coefficient hubs do not overlap strongly with network variants, areas of extreme individual deviation in functional connectivity. However, it is possible that more subtle forms of variation would be present at these hubs that are not captured by the extreme network variants. To investigate this question, we examined continuous measures of similarity at participation coefficient hubs (see Section 2.6.2).

We measured the individual-to-group similarity of whole-brain functional connectivity for the top 10 participation coefficient hubs (Fig. 6). Participation coefficient hub locations generally had good spatial correlations with the group-average connectivity profile in both HCP (r = .59+/-.04) and MSC participants (r = .64+/- .04). Although the pattern of hub connectivity is generally consistent across subjects, there are some deviant cases like MSC09 hub 3 which will be explored further in sections 3.5 and 3.6). Comparisons with random rotations confirmed that hub regions show similar correspondence to the group average as other regions of cortex (see Supp. Fig. 3 and Supp. Table 1; all *p*(FDR) > .74). This suggests that participation coefficient hubs do not differ substantially across individuals, even in more subtle forms of variation.

**Figure 6:**
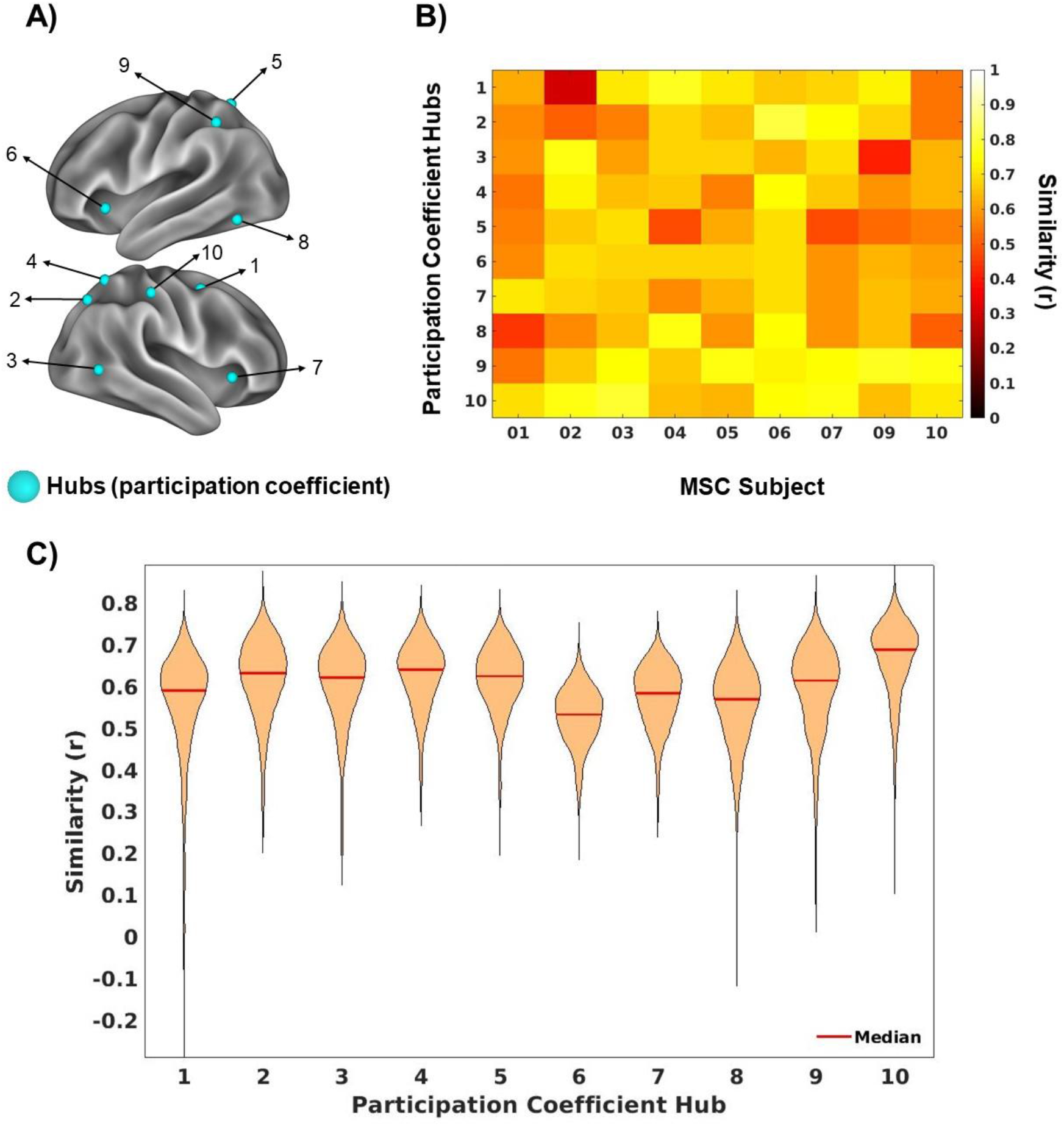
Continuous measures of individual-to-group similarity for participation coefficient hub locations. A) The location of group-average participation coefficient hubs are represented by the light blue circles, numbered for comparison to panels B and C. B) Each column of the x axis represents one of the analyzed MSC subjects and each row corresponds to one of the top participation coefficient hubs (corresponding number). The color scale represents the similarity of FC for a hub in each MSC individual relative to the group average, with warmer colors representing greater similarity. For the most part hubs are similar to the group average connectivity profile. (C) The same measures were calculated for the 752 HCP participants and are represented in a violin plot. The median similarity is marked by a red line.

### 3.5 Characterizing hubs based on their cross-network profiles

The current findings suggest that hubs are largely similar in whole-brain functional connectivity across individuals, in the bounds of what would be expected for other regions of cortex. Next we examined which networks each participation coefficient hub was connected with in individuals, to formally address the conserved vs. malleable hub hypotheses posed in the introduction. For each hub in each MSC participant, we measured its connectivity to 14 canonical networks, creating a network profile for that region.

These profiles are shown for the top 10 participation coefficient hubs in Figure 7. Many hubs showed strong connectivity between 2 networks (1, 2, 5, 8), while others appeared to connect with a broader set (e.g., hub 3, 10). The network profiles were generally consistent across the MSC participants and the group average, in accord with the conserved hub hypothesis. However, some exceptions were present (e.g., MSC02 for Hub 1).

**Figure 7:**
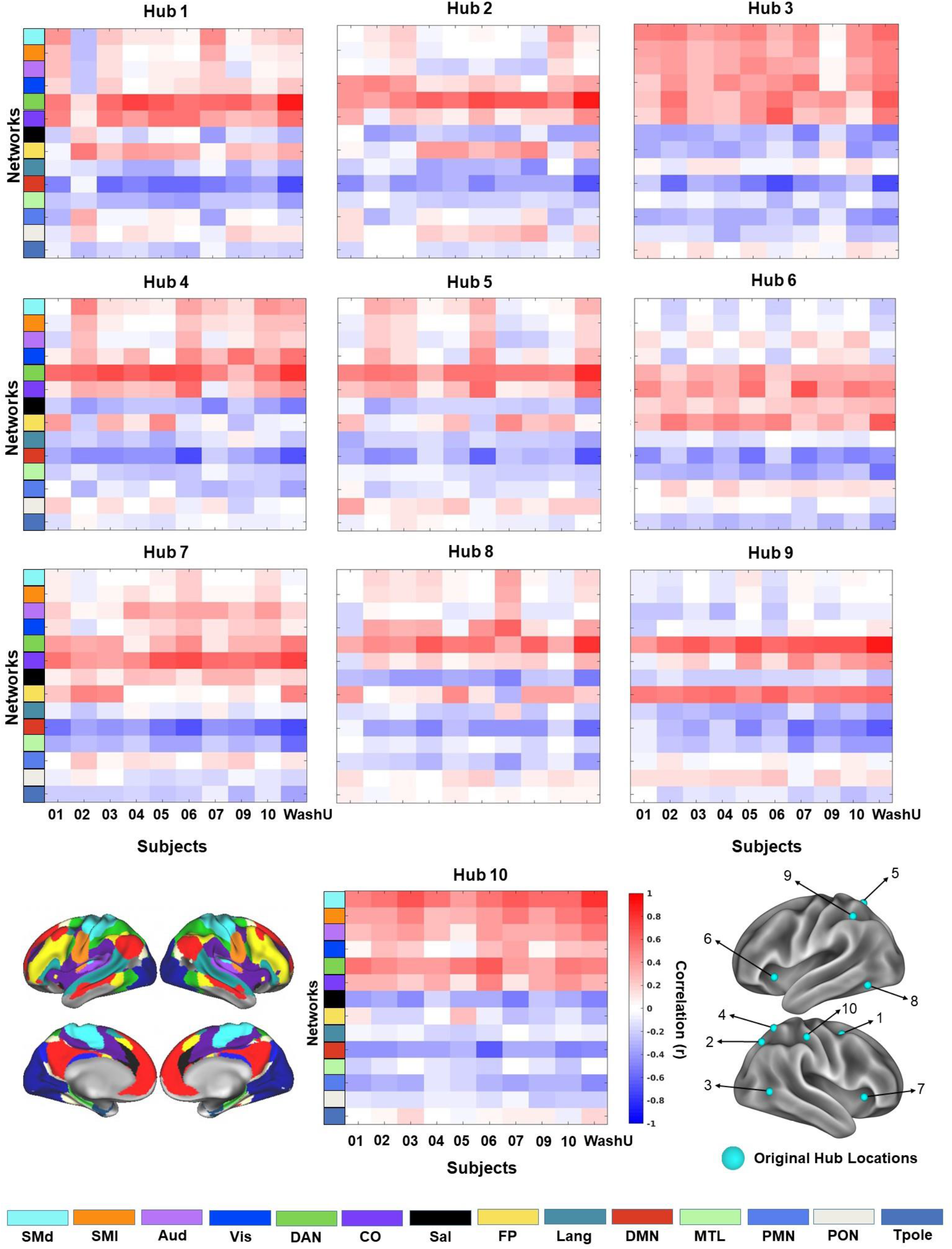
Network profiles for group-average participation coefficient hubs. For each hub (color map sub-plots), we show the network profile for the 9 MSC participants (columns) and WashU-120 group average (final column) to 14 canonical networks (rows; see *Methods*). The hub locations are shown in the bottom right corner and the canonical network maps are shown in the bottom left corner (each network is represented by a color).

### 3.6 Hub locations improve in correspondence if adjusted slightly in position

We observed some exceptions to the conservation in network profiles (Fig. 7) and similarity (Fig. 6) across participants (e.g., MSC02 for Hub 1, MSC09 for Hub 3). We next asked whether these exceptions could be ameliorated if hub locations were allowed to shift slightly in location between individuals. In order to investigate this possibility, we developed a spotlight procedure, in which hubs were adjusted slightly in position within each individual within a 10mm radius to find the location with the best matching network connectivity profile (see *Methods; Fig. 8*).

**Figure 8:**
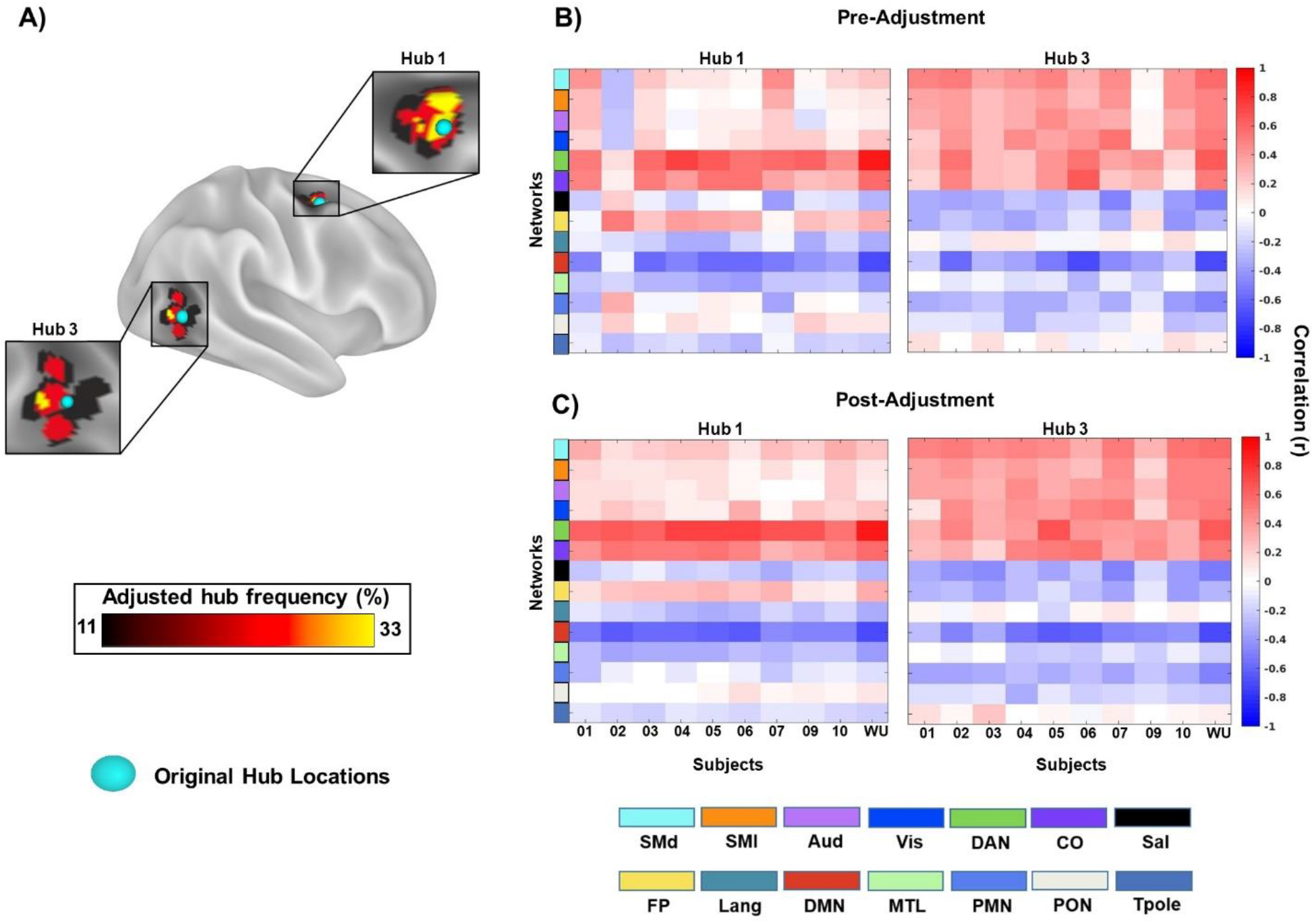
Hub local adjustment procedure. Results of a spotlight search procedure for two representative example hubs (Hub 1 and Hub 3) show that for most subjects hub like areas can be found within a tight zone around a group hub. A) The blue spheres represent the original locations for group hubs 1 and 3. The underlying color map depicts the final adjusted hub locations across the 9 individuals in the MSC (see Supp. Fig 3 for adjusted hub locations for all hubs). (B) The original network profiles are shown for hub 1 (left) and hub 3 (right). (C) The adjusted hub network profiles are shown for the same two hubs. Adjusting hub locations improved correspondence across participants, especially in exception cases (e.g., MSC02 for hub 1, MSC09 for hub 3).

In most cases regions can be found in the spotlight that have a fair degree of resemblance to the group-average network profile (see two representative example hubs in Figure 8). In particular, note that the subjects that were previously exceptions to the general pattern (MSC02 for hub 1, MSC09 for hub 3) show higher correspondence with the group average network profile at a nearby location.

The adjusted hub network profiles are shown for all hubs below in Figure 9. In comparing Figure 7 to Figure 9, note the enhanced consistency in network profile for hubs across individual participants. This enhanced correspondence argues for group hubs being conserved across individuals, if you allow for a relatively minor local adjustment in location.

**Figure 9:**
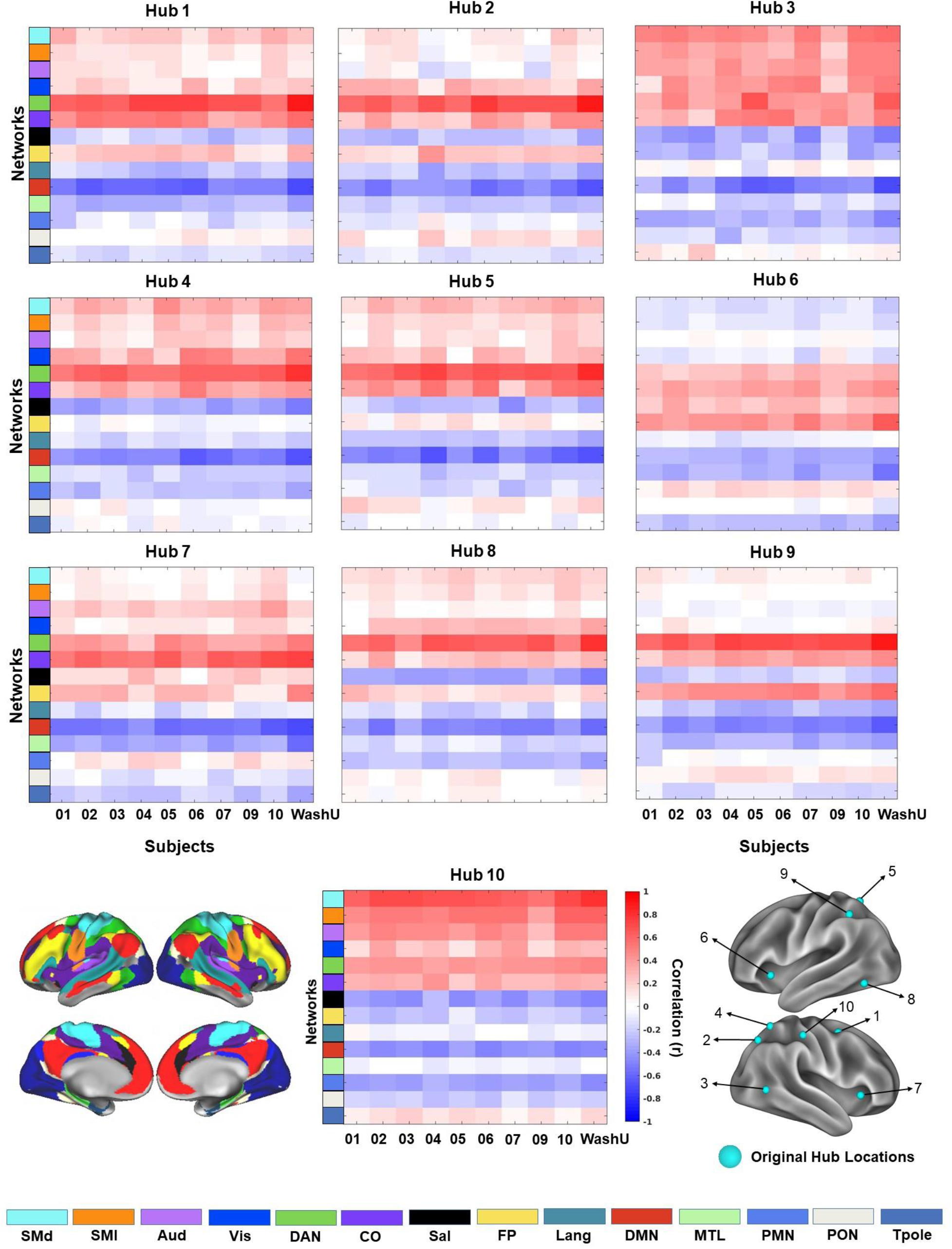
Network profiles for group hubs after local adjustment. This figure is similar to Figure 7, but shows the network profiles for each of the 10 top participation coefficient hubs after those hubs have been slightly adjusted in location using our spotlight analysis procedure (see *Methods* and Supp. Fig. 3 for final adjusted locations). For each hub, we show the network profile for the 9 MSC participants (columns) and WashU-120 group average (final column) to 14 canonical networks (rows). The original hub locations are reproduced in the inset on the lower left, and canonical networks are shown in the lower right.

With this enhanced correspondence, it is also clearer that adjusted hubs seem to fall into one of two patterns. The first pattern is marked by the hub exhibiting a high degree of similarity with two to three networks. This pattern characterizes hubs 1, 2, 5, 8, and 9 (see Fig 9C & Fig 10). For example, hub 1 primarily bridges the dorsal attention and cinguloopercular networks, with weaker links to the frontoparietal and somatomotor systems. The other pattern is a high degree of similarity with a wide set of networks (see Fig 9C & Fig 10). For example, Hub 3 shows strong links to a range of sensorimotor and control networks.

## 4: Discussion

Our goal in this work was to determine how group hubs vary across individuals. We hypothesized that hubs, which are thought to be regions critical to many important functions, should be relatively conserved across individuals. However, we also considered the alternatives that (1) hubs, as regions with diverse connectivity across networks may also show malleability across subjects or (2) that hubs (at the group level) may be artifactual, driven by variation in functional networks within single individuals both of which would be associated with high correspondence between inter-subject connectivity profile variation and connector hub location. We demonstrated that group hubs, defined with the participation coefficient, do not overlap with locations of idiosyncratic functional connectivity and show relatively good correspondence in functional connectivity and network profiles across participants, with further improvements observed if these hubs are adjusted slightly in location per individual. These findings are most consistent with the idea that participation coefficient hubs are relatively conserved across individuals, rather than artifactual or malleable. More caution is warranted with alternative hub measures, such as the proximity based community density metric, which did tend to overlap areas of idiosyncratic functional connectivity. Overall, the findings lend support to the idea that group hubs have relatively consistent functional connectivity characteristics across people, rather than representing sites of particular malleability or artifacts.

### 4.1 Group connector hubs are relatively conserved across people

The participation coefficient is a measure of connection diversity, measuring the extent to which a given node shows connections across multiple distributed networks. Many investigators (Bertolero et al., 2018; Bertolero et al., 2015; Cole et al., 2013; Gordon et al., 2018; Gratton et al., 2016; Gratton et al., 2012; Power et al., 2013; Warren et al., 2014) have identified brain connector hubs using this metric, often based on group-average data (Power et al., 2013). Although these regions are defined by their diverse connection profile and are located in association regions of the brain (which show high inter-subject variation; Kong et al., 2019; Mueller et al., 2013; Seitzman et al., 2019), here we show that they do not have a strong correspondence to diversity *across people:* that is, diversity in functional connectivity across networks does not track with diversity in connectivity across individuals. Group-average hubs defined with the participation coefficient do not frequently overlap with extreme variants (Fig 3), are relatively similar to the group average (Fig 7), and have a fairly consistent network profile (Fig. 8; especially if these hubs are allowed to move slightly in position across people; see Fig 9–10). The lack of correspondence between group hubs and inter-subject connectivity variation was observed in two datasets and was true for both extreme deviations from the group average connectivity profile (variants) and more subtle deviations (similarity to the group).

This relative lack of variation is consistent with previous literature, which has shown that group-average estimates of hubs serve as good priors for locations that have outsized impacts on network structure and behavior if damaged in lesion patients (Gratton et al., 2012; Warren et al., 2014). We and others have also proposed that connector hubs (defined by the participation coefficient) play a critical role during task execution (Bertolero et al., 2018; Cole et al., 2013; Gratton et al., 2016). Hub regions are activated across a range of cognitive processes (Bertolero et al., 2015) and hub connectivity is modulated by task context (Cole et al., 2013; Gratton et al., 2016). Moreover, some evidence shows that participation coefficient hubs promote network modularity during task performance by tuning the connectivity of their neighbors (Bertolero et al., 2018), This body of work suggests that hubs may help coordinate activity between networks as needed in task control. This is a relatively essential set of functions to everyday life, likely depending on the ability to enact specific patterns of cross-network connectivity, and thus requiring some degree of uniformity in connector hub profiles across subjects.

However, although not *more* variable than would be expected by chance, participation coefficient hubs were also not generally below the expected variation for the cortex. The lack of a significant difference was observed across two datasets (HCP and MSC) and across multiple investigative approaches focusing on both extreme deviations and continuous gauges of similarity. Thus, group hubs may be relatively conserved, but not more so than many other regions of the brain. Therefore, while these results argue that group (participation coefficient) hubs are not especially malleable or driven by artifacts, it is likely not correct to interpret them as particularly more conserved than other cortical regions. Indeed, although recent work has highlighted strong variation between individuals (Bijsterbosch et al., 2018; Finn et al., 2015; Gordon et al., 2017a; Gratton et al., 2018a; Kong et al., 2019; Miranda-Dominguez et al., 2014; Mueller et al., 2013), much of this variation is relatively punctate and restricted to particular locations in a particular person (Seitzman et al., 2019), leading many places in the cortex to show good correspondence to the group average (Seitzman et al., 2019). It is possible that improved individual measures of hubs will identify regions with better conservation (e.g., see section 3.6 – Fig 10 and section 3.7 – Fig 11).

### 4.2 Individual measures of brain hubs may bring added precision

We have shown that hubs defined with group-average data have connectivity profiles that are relatively conserved. Yet, it is still not clear if these group-average hubs identify the best hubs within single individuals. The results of our spotlight analysis (Fig. 9–10) suggest that, at a minimum, researchers should consider exploring the areas surrounding group-average hub locations to ameliorate correspondence across participants.

Even when group hubs are reproducible across individuals, they could miss important features of functional neuroanatomy that can only be observed when looking at individually-defined hubs. It might be the case that group hubs detect hub locations that are conserved across individuals, but that other hubs exist within each person that are more malleable and difficult to capture in group maps. Hubs defined in single individuals may also contain further information that is obscured in group hubs. For example, individually-defined hubs might be needed to uncover hub sub-types, which may help uncover more refined properties associated with cognition (Gordon et al., 2018).

Procedures that adopt individual-level connectivity maps, like the spotlight analysis we employed, could be further refined to help answer these questions. The high reliability of precision fMRI data makes it especially well-suited to ensuring robust applications of these procedures (Gordon et al., 2017b). In addition, high resolution imaging made possible by 7T scanners would allow for greater confidence in the accuracy of individual-level maps through improved signal and removal of artifactual hubs caused by spatial blurring or close proximity (Braga et al., 2019; Viessmann and Polimeni, 2021). Projects using these techniques may help to further chart correspondence between the location of hubs defined on individual subject data and hubs defined at the group level, which do not always show very tight correspondence (Gordon et al., 2018).

### 4.3 Variants are not strong hubs

Thus far, we have framed our results and discussion in terms of what our findings reveal for hubs. However, our findings also provide insights into variant locations. As locations of idiosyncratic functional connectivity, it is possible that variants are manifestations of flexible bridges between networks, which drives their inconsistency in their network membership across subjects (Vázquez-Rodríguez et al., 2019; Zhang et al., 2016). Yet, we did not find a clear correspondence between variants and participation coefficient hubs across several analyses in two datasets. While there is a correspondence between variants and community density hubs, this may be artifactual, due to averaging across people, rather than a manifestation of variants arising from a hub-like nature. These results suggest that, as a general rule, variants do not overlap with sites of group level connector hubs, implying that variants do not have a functional role as chameleon-like integrative regions. However, it is still possible that specific variants may show overlap with individual-level hubs that are not captured with this group approach. Future work will be needed to further investigate the properties of variants and their correspondence to individual-level hubs.

### 4.4 Community density is a hub metric that should be used with caution in the group

Unlike hubs defined by the participation coefficient, community density hubs defined in group-average data had a relatively strong correspondence with idiosyncratic variant regions. Community density hubs are defined based on their proximity to multiple different networks (Power et al., 2013), following the assumption that hubs should be positioned spatially intermediate to the systems that they bridge. However, although distributed somewhat similarly to participation coefficient hubs (Power et al., 2013), community density hubs in the group average are found more ventrally in the anterior insula, along the superior frontal cortex, and near the temporoparietal junction – locations in closer accord with locations of inter-subject variation (Fig 5).

This correspondence with regions of inter-subject variation suggests that group hubs defined with community density represent either malleable regions or artifactual hubs caused by mixed signals across individuals. Individual differences in brain networks often occur near the boundaries between systems (Seitzman et al., 2019). As community density hubs sit at a nexus of multiple networks, then they may be more vulnerable to contamination from these shifting boundaries and the appearance of artifactual hubs. Future work using community density hubs should be cautious of this potential correspondence, especially if the hubs are identified with group average data. A solution to this issue may be to identify community density hubs within individual subject data instead, where hub inflation caused by inter-subject variation can be ruled out. In those cases, it may be possible to identify better representations of community density hubs that help to mediate interactions between networks based on their proximity.

### 4.5 Conclusion

Our findings from two independent datasets suggest that group-average brain hubs are relatively consistent in their functional topography across individuals. These findings suggest that hub locations may be a relatively conserved property of brain networks, perhaps due to the critical role these regions have been proposed to have in cognition and brain function. Alternative metrics for hubs based on spatial proximity, such as community density, can overlap more strongly with locations of individual variability and should be used with more caution in the group average.

## Acknowledgments

This work was supported by an NSF CAREER 2048066 (CG) and NIH grants R01MH118370 (CG), T32NS047987 (DMS, BTK). This research was supported in part through the computational resources and staff contributions provided for the Quest high performance computing facility at Northwestern University which is jointly supported by the Office of the Provost, the Office for Research, and Northwestern University Information Technology.

## Footnotes

1 Note that there are many ways to define hubs within complex systems, ranging from classic degreebased measures, to measures of centrality, to connector hubs (Bullmore, & Sporns, 2012; Van Den Heuvel, & Sporns, 2011). For the purposes of this work, we focus on connector hubs: because of their links to different networks, connector hubs are well situated for transferring information across functional systems and have been the focus of several past studies on the importance of hubs in the human brain (Bertolero et al., 2015; Cole et al., 2013; Gratton et al., 2012; Gratton et al., 2016; Power et al., 2013).

