## Supplemental Material for "Brain hubs defined in the group do not overlap with regions of high inter-individual variability"

#### Supplemental Tables

#### Table S1: Similarity to the Group for MSC Subjects

| MSC subject | Mean *R* to group | 95% CI |
|---|---|---|
| MSC01 | .57 | [0.47, 0.65] |
| MSC02 | .63 | [0.56, 0.73] |
| MSC03 | .67 | [0.53, 0.68] |
| MSC04 | .66 | [0.56, 0.70] |
| MSC05 | .65 | [0.55, 0.71] |
| MSC06 | .71 | [0.57, 0.75] |
| MSC07 | .65 | [0.55, 0.72] |
| MSC09 | .62 | [0.52, 0.66] |
| MSC10 | .61 | [0.57, 0.72] |

Supp. Table 1: For each MSC subject the mean correlation (across vertices included in the hub) is shown in the middle column. The third column contains the corresponding 95% confidence intervals.

### Supplemental Figures

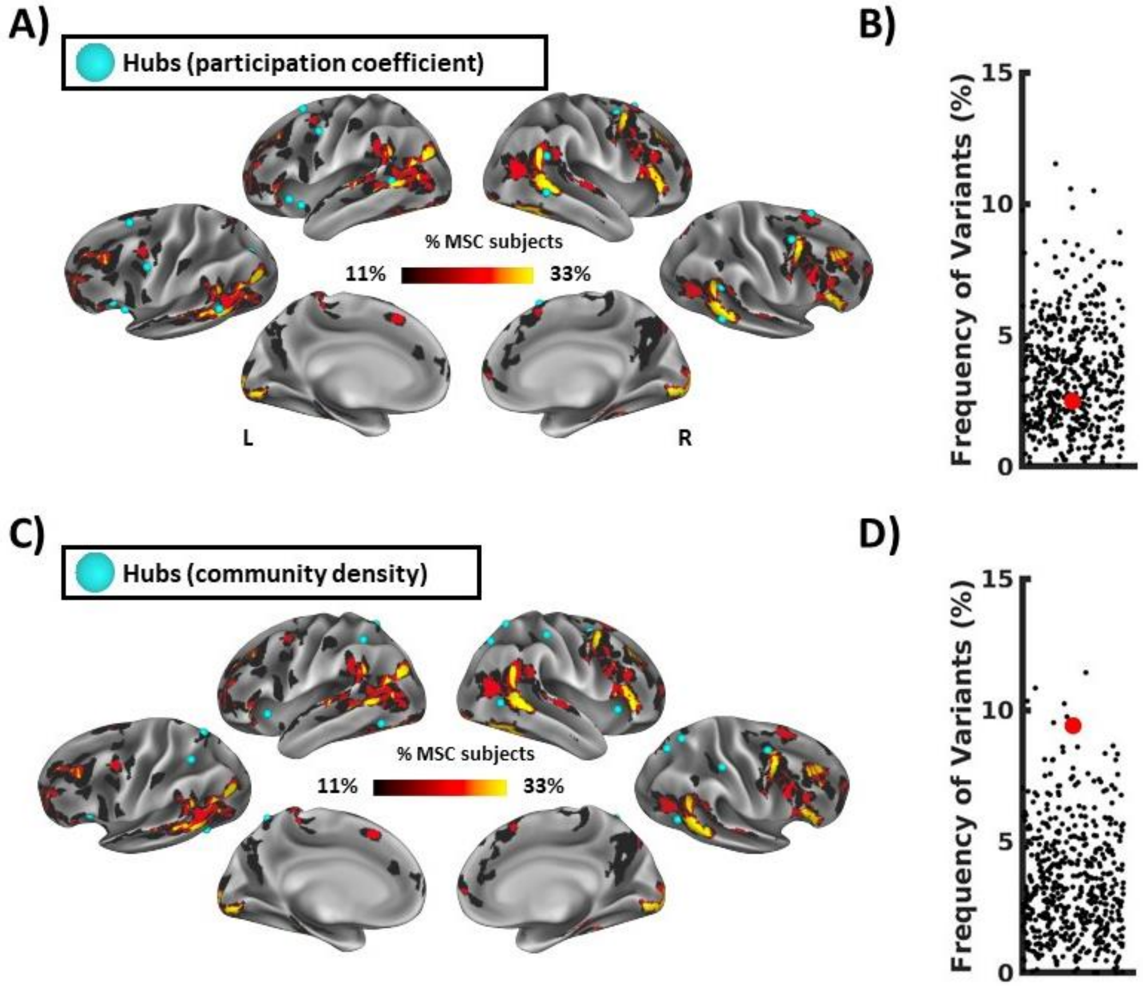


Figure S1: Locations of group hubs relative to the frequency of network variants in the Midnight Scan Club (MSC) dataset. A) This figure plots the locations of the participation coefficient hubs (light blue foci) overlapped on frequency of network variants in the MSC dataset. B) This figure displays the frequency of networks variants at participation coefficient hubs (red dot) relative to random rotations (black dots). In the MSC, 2.49% of people had network variants at participation coefficient hubs (36$^{th}$ percentile of rotations, 95% CI [0.38%, 8.09%]). C) This figure is similar to A, but shows locations of community density hubs overlapped on the frequency of network variants in the MSC dataset. D) As in B, this figure shows the frequency of network variants at community density hubs (red dot) relative to random rotations (black dots). In the MSC, 9.40% of people had network variants at community density hub locations, corresponding to the 99$^{th}$ percentile of random rotations (95% CI [0.29%, 7.95%]).

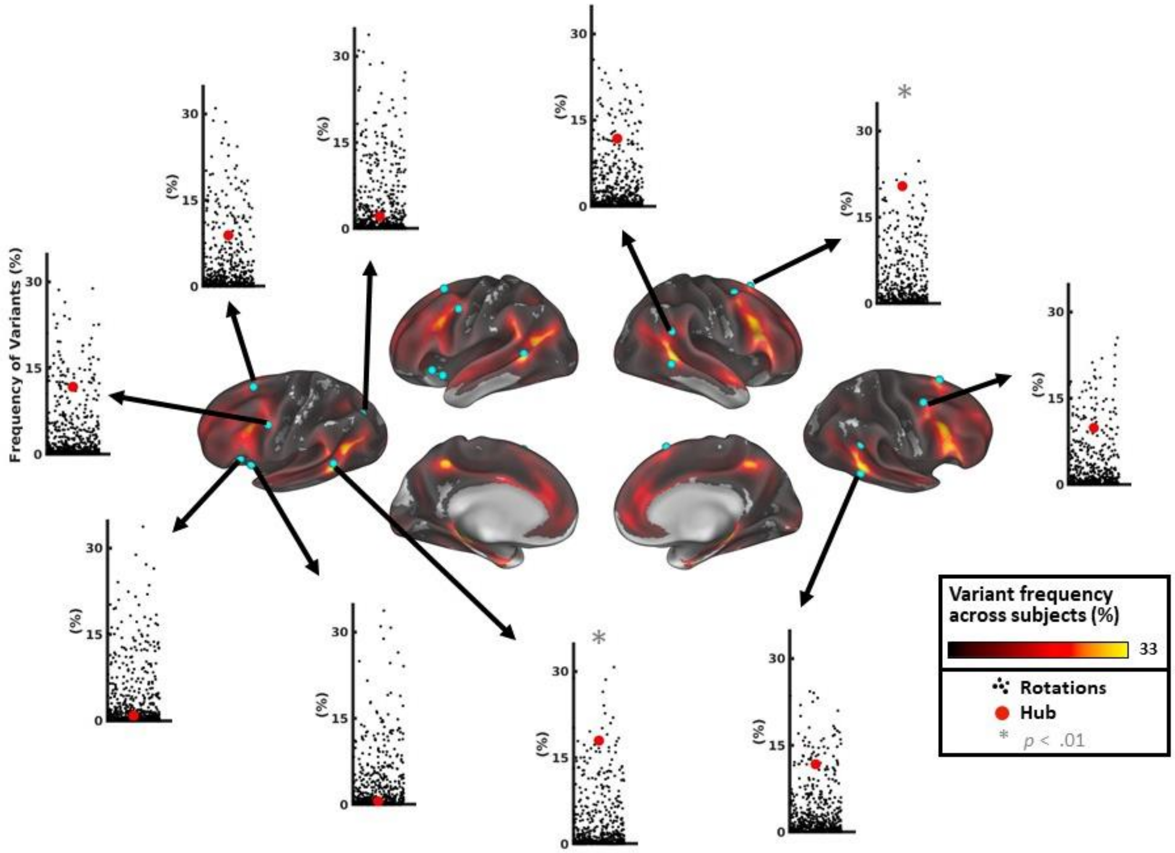


Figure S2: Network variant frequency at single community density hub locations. Single community density hub locations tend to exhibit a high degree of variant overlap. Each foci on the cortical surface represents a community density based hub. The scatter plots display the relationship between true frequency of variants in a hub region (red dot) and the amount expected by random rotations (black dots). All but three hubs fell above the mean of their respective hemisphere's null distribution. An asterisk placed on the scatter plot denotes hubs that significantly differed from what would be expected by the random rotations (* p<0.01), but no effects survived FDR correction.

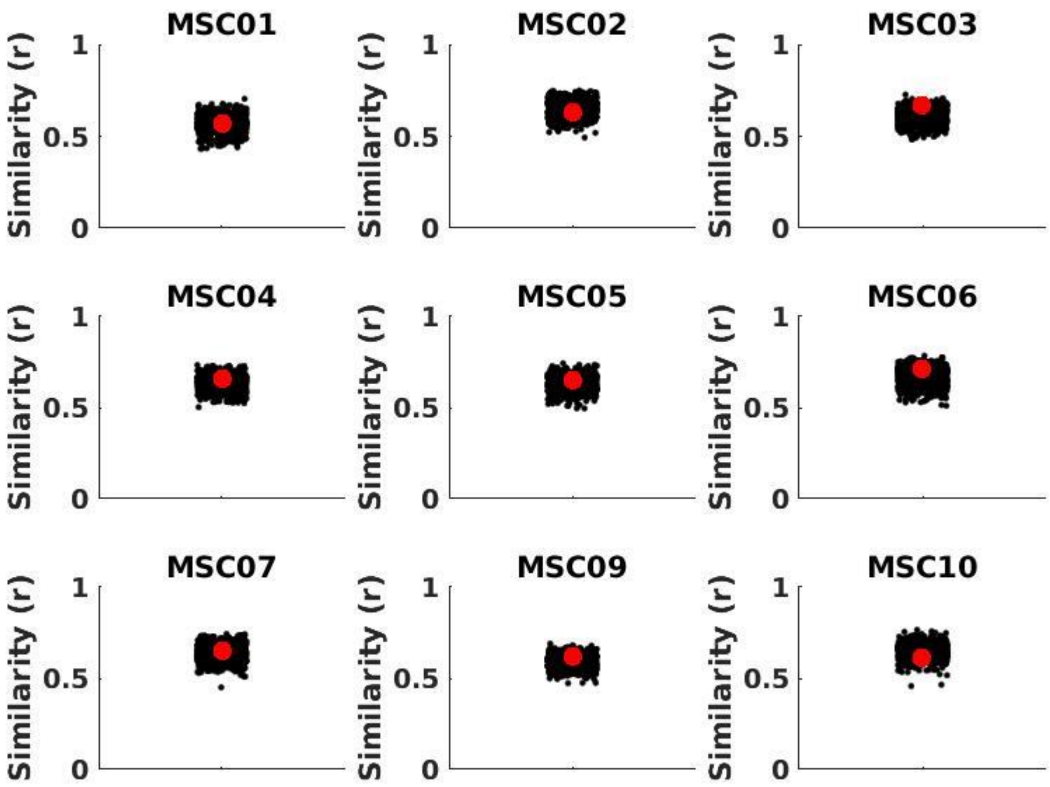


Figure S3: Similarity to the group average connectivity profile distributions for each MSC subject. For each MSC participant the similarity of hub locations to the group average connectivity profile is represented by the red dot and the smaller black dots represent the similarity for random rotations of the hub set. None of the participants significantly deviated from what would be expected by the null distribution (see Supp. Table 1 for confidence intervals). The average observed similarity to the group connectivity profile (average of the red dots) across participants was .64 with a standard deviation of .04.

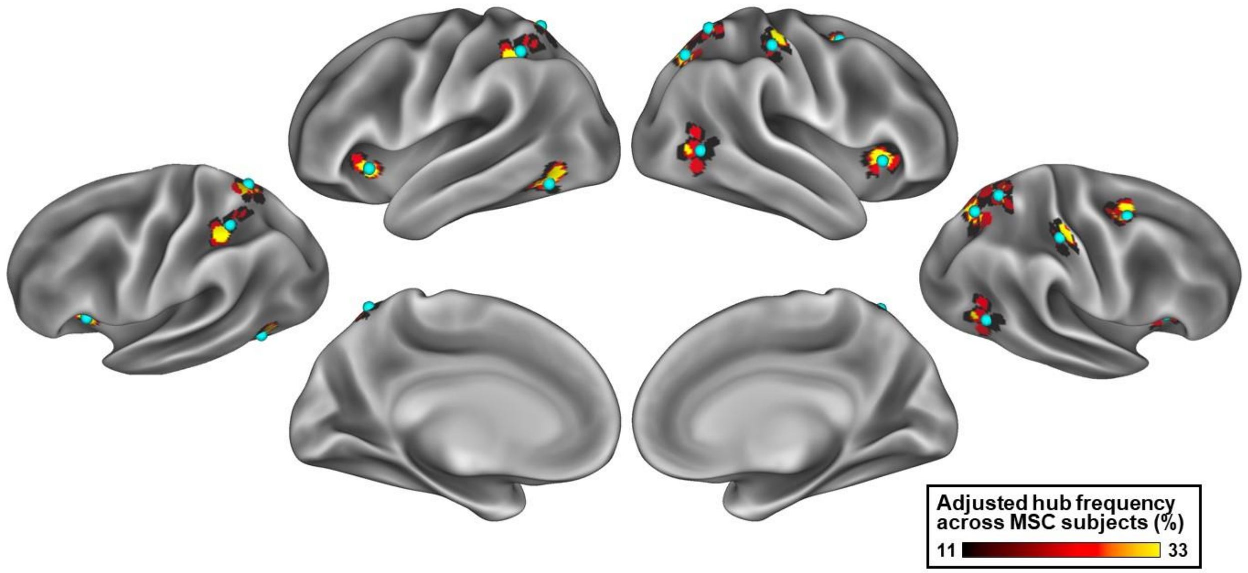


Figure S4: For each group-average participation coefficient hub (blue spheres) the frequency of adjusted hub locations is marked. The frequency of adjusted hub locations is defined as the percentage of MSC subjects with an adjusted hub overlapping with the given cortical location.
